# Glycogen degrading activities of catalytic domains of α-amylase and α-amylase-pullulanase enzymes conserved in *Gardnerella* spp. from the vaginal microbiome

**DOI:** 10.1101/2022.10.19.512974

**Authors:** Pashupati Bhandari, Jeffrey Tingley, D. Wade Abbott, Janet E. Hill

## Abstract

*Gardnerella* spp. are associated with bacterial vaginosis, in which normally dominant lactobacilli are replaced with facultative and anaerobic bacteria including *Gardnerella* spp. Co-occurrence of multiple species of *Gardnerella* is common in the vagina and competition for nutrients such as glycogen likely contributes to the differential abundances of *Gardnerella* spp. Glycogen must be digested into smaller components for uptake; a process that depends on the combined action of glycogen degrading enzymes. In this study, the ability of culture supernatants of 15 isolates of *Gardnerella* spp. to produce glucose, maltose, maltotriose and maltotetraose from glycogen was demonstrated. Carbohydrate active enzymes were identified bioinformatically in *Gardnerella* proteomes using dbCAN2. Identified proteins included a single domain α-amylase (EC 3.2.1.1) (encoded by all 15 isolates) and an α-amylase-pullulanase (EC 3.2.1.41) containing amylase, carbohydrate binding modules and pullulanase domains (14/15 isolates). To verify the sequence-based functional predictions, the amylase and pullulanase domains of the α-amylase-pullulanase, and the single domain α-amylase were each produced in *E. coli*. The α-amylase domain from the α-amylase-pullulanase released maltose, maltotriose and maltotetraose from glycogen, and the pullulanase domain released maltotriose from pullulan and maltose from glycogen, demonstrating that the *Gardnerella* α-amylase-pullulanase is capable of hydrolyzing α-1,4 and α-1,6 glycosidic bonds. Similarly, the single domain α-amylase protein also produced maltose, maltotriose and maltotetraose from glycogen. Our findings show that *Gardnerella* spp. produce extracellular amylase enzymes as ‘public goods’ that can digest glycogen into maltose, maltotriose and maltotetraose that can be used by the vaginal microbiota.

**Importance:** Increased abundance of *Gardnerella* spp. is a diagnostic characteristic of bacterial vaginosis, an imbalance in the human vaginal microbiome associated with troubling symptoms, and negative reproductive health outcomes, including increased transmission of sexually transmitted infections and preterm birth. Competition for nutrients is likely an important factor in causing dramatic shifts in the vaginal microbial community but little is known about the contribution of bacterial enzymes to the metabolism of glycogen, a major food source available to vaginal bacteria. The significance of our research is characterizing the activity of enzymes conserved in *Gardnerella* species that contribute to the ability of these bacteria to utilize glycogen.

## Introduction

*Gardnerella* spp. are associated with bacterial vaginosis (BV), a condition characterized by the replacement of the usually dominant *Lactobacillus* community with a mixture of facultative and anaerobic bacteria including *Gardnerella* spp. (1). BV is characterized by thin, white to greyish discharge and unpleased ‘fishy’ vaginal odor, however a large majority of women diagnosed with BV do not have symptoms (2). Phenotypic diversity among *Gardnerella* spp. may explain why they are found in women presenting clinical signs of bacterial vaginosis and in asymptomatic or BV negative women (3). The genus *Gardnerella* has been classified into four different species and at least 13 “genome species” (4). Co-occurrence of multiple *Gardnerella* species in the vaginal environment is common, and different species are dominant in different women (5). Given the potentially different roles of *Gardnerella* species in the vaginal microbiome, understanding the factors contributing to their population structure is important. Previous work has shown that interactions in co-cultures of *Gardnerella* spp. are typical of a “scramble” competition (6). In scramble competition, one competitor outgrows others through its superior ability to use shared resources such as nutrients (7). Competition for nutrients could be an important factor contributing to the differential abundances of *Gardnerella* spp. in the vaginal microbiome.

Glycogen is an abundant carbon and energy source available to the vaginal microbiota including *Gardnerella*. Glycogen is a glucose homopolysaccharide comprising linear a-1,4 linked glucose with α-1,6 branch points occurring approximately at every 8-13 residues to from a highly branched and complex structure. The sizes of glycogen molecules and branching patterns can vary based on the glycogen source (8). In the vagina, glycogen accumulates inside vaginal epithelial cells under the influence of estrogen (9) and is released into the vaginal lumen through the activity of bacterial cytolysins and/or lactic acid mediated acidosis (10). The amount of free glycogen in vaginal fluid can range from undetectable to 32 μg/ml (11).

Glycogen must be digested into smaller products for uptake into bacterial cells. This complex and multistep process requires the combined action of enzymes collectively known as glycogen degrading enzymes. The majority of Glycogen degrading bacterial amylases belong to the glycosyl hydrolase (GH) class within family 13 in the classification of carbohydrate active enzymes. GH13 enzymes are further classified into more than 40 subfamilies based on their structure and substrate specificity (12). Previous studies have shown the presence and activity of human and/or bacterial amylases in vaginal fluid (13, 14) but the specific contribution of *Gardnerella* spp. to glycogen digestion is not clear.

This study aimed to assess the glycogen digestion ability of *Gardnerella* spp. isolated from the vaginal microbiome and to determine the distribution of extracellular glycogen degrading enzymes among different species. In addition, we aimed to biochemically characterize the activities of the predicted functional domains of extracellular glycogen degrading amylase enzymes identified. Our findings showed that *Gardnerella* spp. have α-amylase and α-amylase-pullulanase enzymes which can digest glycogen into maltose and other oligosaccharides.

## Methods

### *Gardnerella* isolates

Isolates of *Gardnerella* used in this study were from a previously described culture collection (15) and included a total of 15 *Gardnerella* spp. isolates [three representative strains each from *G. leopoldii* (GH005, VN003 & NR017), *G. piotii* (VN002, GH007 & GH020), *G. vaginalis* (NR038, NR001 & NR039) and *G. swidsinskii* (NR016, NR021 & NR020), two isolates from *Gardnerella* genome sp. 3 (NR026 & N170) and one from unknown genome species (NR047, subgroup D based on cpn60 classification system). Whole genome sequence data for all isolates were retrieved from NCBI BioProject Accession PRJNA394757.

### Amylase activity assay

Amylase activity of *Gardnerella* isolates was assessed using a glycogen iodine test (16). Isolates were revived from −80°C storage on Columbia blood agar plates containing 5% sheep blood, and incubated anaerobically using the GasPak system (BD, USA) at 37°C for 48 hours. Isolated colonies were transferred into modified NYCIII broth (obtained by replacing horse serum with bovine serum and omitting glucose in NYCIII media) and incubated anaerobically for 24 hours at 37°C. Overnight broth cultures were centrifuged at 10,000 × *g* for 10 mins and the cell-free supernatant was passed through a 0.45 μm filter. Filtered supernatant (10 μL) was spotted on to a 1% (w/v) glycogen (Bovine liver glycogen, Sigma) agar plate and incubated anaerobically at 37°C for 24 hours. Filtered supernatant was also streaked on Columbia sheep blood agar to confirm sterility. Plates were flooded with 1% (w/v) Lugol’s iodine solution to detect evidence of amylase activity. α-amylase from *Bacillus licheniformis* (0.05 mg/ml) (Sigma-Aldrich, Oakville, ON, CAT No. A455) and modified NYCIII broth were used as positive and negative controls respectively.

Activity of cell culture supernatants on glycogen was assessed by incubating culture supernatants with glycogen and examining the products. Briefly, 20 μl of culture supernatant was added to 180 μl of 0.1% bovine liver glycogen in 50 mM sodium phosphate buffer, pH 7.0 and incubated at 37°C. Aliquots (50 μl) of the reaction mixture were collected at 24 h and were frozen immediately until analysis by thin layer chromatography (TLC) and high-performance anion exchange chromatography with pulsed amperometric detection (HPAEC-PAD). TLC was performed on silica plates in a 1-butanol, acetic acid and distilled water (2:1:1 [vol:vol:vol]) mobile phase and stained using 0.3% of N-napthyl ethylenediamine dihydrochloaride (Sigma) as described previously (17). HPAEC-PAD samples were first precipitated with 80% ethanol to remove large polysaccharides and diluted (to 1:20 for G5 and maltodextrin or 1:10 for glycogen) and resolved on a Dionex PA20 column in a mobile phase of 30 mM NaOH and gradient of 10 mM to 120 mM sodium acetate over 40 min at a flow rate of 0.5 ml min^-1^. Data were analyzed using Chromeleon v6.80 chromatography data system software.

### Identification of the carbohydrate active enzymes (CAZyme)

The *Gardnerella* proteomes, predicted from the RAST annotation, were uploaded into the dbCAN2 webserver (18). All three available domain prediction tools (HMMER, eCAMI and DIAMOND) were used to annotate the carbohydrate active domain(s) in protein sequences. The dbCAN2 uses SignalP to predict whether a signal peptide is present or absent in the protein in which each carbohydrate active domain is located. To identify secreted glycogen degrading glycosyl hydrolases, dbCAN2 output was filtered to include only GH13 domain(s) associated with signal peptide. TMHMM 2.0 (19) and LocateP (20) were used to predict transmembrane and cell-wall anchored domains respectively.

### Phylogenetic analysis of GH13_32 amylase domains

Domains belonging to subfamily 32 within the family 13 of glycosyl hydrolases class are referred to as GH13_32 amylase domains. Amino acid sequences corresponding to the GH13_32 amylase domain were extracted from predicted extracellular glycosyl hydrolases of all study isolates and four *Gardnerella* reference strains [(*G. leopoldii*, (CP029984)), *G. piotii* (QJUV00000000), *G. vaginalis* (QJUZ00000000) and *G. swidsinskii* (QJVB00000000)](4). Multiple-sequence alignments were performed using CLUSTALw and a neighbour joining consensus tree was built in Geneious prime version 2022.1.1 (https://www.geneious.com). The final tree was visualized using Figtree (v1.4.4).

To identify the conserved catalytic residues in GH13_32 domains, amino acid sequences of representative GH13_32 domains of *G. leopoldii* NR017 were combined with GH13_32 domain sequences from functionally characterized α-amylases [*Pseudoalteromonas haloplanktis* A23 (CAA41481.1) and *Streptomyces venezuelae* ATCC 15068 (AAB36561.1)] and amylopullulanase *[Bifidobacterium breve* UCC2003 (AAY89038.1)] deposited in the CAZy database. Sequences were aligned using CLUSTALw and alignments were visualized in ESPript (21).

### Expression and purification of α-amylase and pullulanase domains from α-amylase-pullulanase

To characterize activities of the amylase and pullulanase domains of the α-amylase-pullulanase, *G. leopoldii* (NR017) was chosen as a representative isolate. Genomic DNA was extracted from *G. leopoldii* NR017 using a modified salting out procedure. Primers (Supplementary Table 1) were designed to amplify the nucleotide sequences encoding the predicted α-amylase (corresponding to amino acids 74-350 of Genbank Accession RFT33565) and pullulanase (corresponding to amino acids 1141-1462 of Genbank Accession RFT33565) domains of the NR017 α-amylase-pullulanase gene along with 30 additional flanking nucleotides each on the 5’ and 3’ ends. PCR was carried out in PCR buffer (0.2 M Tris-HCl [pH 8.4], 0.5 M KCl), a 200 mM concentration of each deoxynucleoside triphosphate (dNTP), 2.5 mM MgCl_2_, a 400 nM concentration of each primer (a-amylase: JH0810-F & JH0811-R, or pullulanase: JH0812-F & JH0813-R), 1 U/reaction of Platinum Taq DNA polymerase high-fidelity (5 U/ml in 50% glycerol, 20 mM Tris-HCl, 40 mM NaCl, 0.1 mM EDTA, and stabilizers) (Life Technologies), and 2 μl of template DNA, in a final volume of 50 μl. PCR was performed with the following parameters: initial denaturation at 94°C for 3 min, 40 cycles of denaturation at 94°C for 15 s, annealing at 60°C for 15 s, and extension at 72°C for 1 min (for α-amylase domain) or 1.5 min (for pullulanase domain), and final extension at 72°C for 3 min. Purified PCR products were digested with *BamHI* and *KpnI* and ligated into expression vector pQE-80L treated with the same restriction endonucleases. The pQE-80L+amylase domain plasmid (henceforth referred to as pQE80L-AMY) was used to transform chemically competent *E. coli* Rosetta DE3 cells (Novagen, Sigma-Aldrich), and the pQE-80L+pullulanase domain plasmid (henceforth referred to as pQE80L-PULL) was used to transform One Shot TOP10 chemically competent *E. coli* cells (Invitrogen, Carlsbad, CA). Insertion of amylase and pullulanase domain encoding sequences in frame with the N-terminal His-tag was confirmed by sequencing of the purified plasmid.

*E. coli* Rosetta cells containing pQE80L-AMY were grown in 800 ml of LB broth containing ampicillin (100 μg/ml) and chloramphenicol (25 μg/ml) whereas *E. coli* Top10 cells containing pQE80L-PULL were grown in same volume of LB broth with ampicillin (100 μg/ml) only. Once the culture reached an optical density at 600 nm (OD_600_) of 0.4 −0.6, protein expression was induced with 0.5 mM IPTG. After 4 h of incubation at 37°C with shaking at 200 rpm, cells were harvested by centrifugation at 8500 × *g* for 20 min at 4°C and pellets were stored at −20°C until further processing. Proteins were solubilized using anionic detergent as described previously (22) with a slight modification during sonication (6 min total run time, 2 min on, 1 min off at 4°C). Eluted proteins were dialyzed against 1 × PBS (pH 7.4) at 4°C using a semipermeable membrane with cut-off size 12 kDa for 24 −48 hours. The diluted proteins were concentrated using a 30 kDa molecular weight cut-off protein concentrator (Thermo Fisher Scientific). Final protein concentration was measured using a Nanodrop spectrophotometer before storing at −20°C.

### Expression and purification of single amylase domain containing α-amylase

A codon-optimized DNA construct corresponding to the full-length open reading frame of single domain α-amylase from *G. leopoldii* NR017 (RFT33564.1) was synthesized and a region encoding amino acids 58-519 (excluding N-terminal signal peptide and C-terminal transmembrane domain) was amplified using JH0857 and JH0858 primers (Supplementary Table 1). The 1463 bp product was ligated into *MfeI and HindIII* digested pET41a+ vector to express as a N-terminal GST-His tag fusion protein (79.9 kDa). The recombinant plasmid (pET41a+CO-AMY) was used to transform chemically competent *E. coli* DH5a cells. Presence of the insert in frame with the N-terminal GST-His-tag was confirmed by Sanger sequencing of the purified plasmid.

Purified recombinant plasmid (pET41a+CO-AMY) was used to transform *E. coli* BL21DE3 cells for protein production. *E. coli* BL21DE3 cells containing recombinant plasmid were grown in 1.8 L of LB broth containing kanamycin (30 μg/ml). When culture reached to optical density OD_600_ 0.4 −0.6, IPTG was added to a final concentration of 0.5 mM to induce protein expression at 37°C for 4 hours with shaking at 200 rpm. Cells were harvested by centrifugation at 8500 × *g* for 20 min. Expressed protein was solubilized using anionic detergent as described previously (22) with slight modification of the sonication protocol (80% duty cycle, 8 minutes total run time, 10 secs on, 5 secs off). The recombinant amylase protein was purified using a nickel column. Eluted proteins were dialyzed against 1× PBS (pH 7.4) at 4°C overnight using a semipermeable membrane of cut-off size 12 kDa, and dialyzed protein was concentrated using a 30 kDa molecular weight cut-off protein concentrator (Thermo Fisher Scientific). Final protein concentration was measured using a Nanodrop spectrophotometer before storing at −20°C.

The region encoding the GST and 6xHis tags was expressed from empty pET41a vector in *E. coli* BL21DE3 cells and was purified using Ni-affinity chromatography for use as a negative control in activity experiments.

### Enzyme activity on different substrates

The activity of purified enzymes was tested by incubating purified enzymes with different substrates. Briefly, 250 μl of the purified amylase domain (22.3 μM) or α-amylase (13.0 μM) was incubated with 1% glycogen, maltodextrin (dextrose equivalent 13-17) or maltopentose while purified pullulanase (25.3 μM) was incubated with 1% maltotriose, maltotetraose, maltopentose, pullulan and glycogen. To test the combined activity of α-amylase and pullulanase domains, an equimolar (25.3 μM) mixtures of both enzymes were incubated with glycogen (1%). All substrates were prepared in 50 mM sodium phosphate buffer at pH 6.0. Enzyme substrate reaction mixtures were incubated at 37°C for 24 h and stored at −20°C until further analysis by TLC or HPAEC-PAD as described above.

## Results

### Amylase activity of *Gardnerella* culture supernatants

No growth was observed from filtered culture supernatants, confirming their sterility. Supernatants of all 15 isolates showed amylase activity in the agar plate assay (Figure 1). Cell-free culture supernatants of 14 isolates produced glucose, maltose, maltotriose and maltotetraose from glycogen while glucose was the only product detected from N170 supernatant (*Gardnerella* genome sp. 3, GS3-2) (Figure 2).

**Figure 1:**
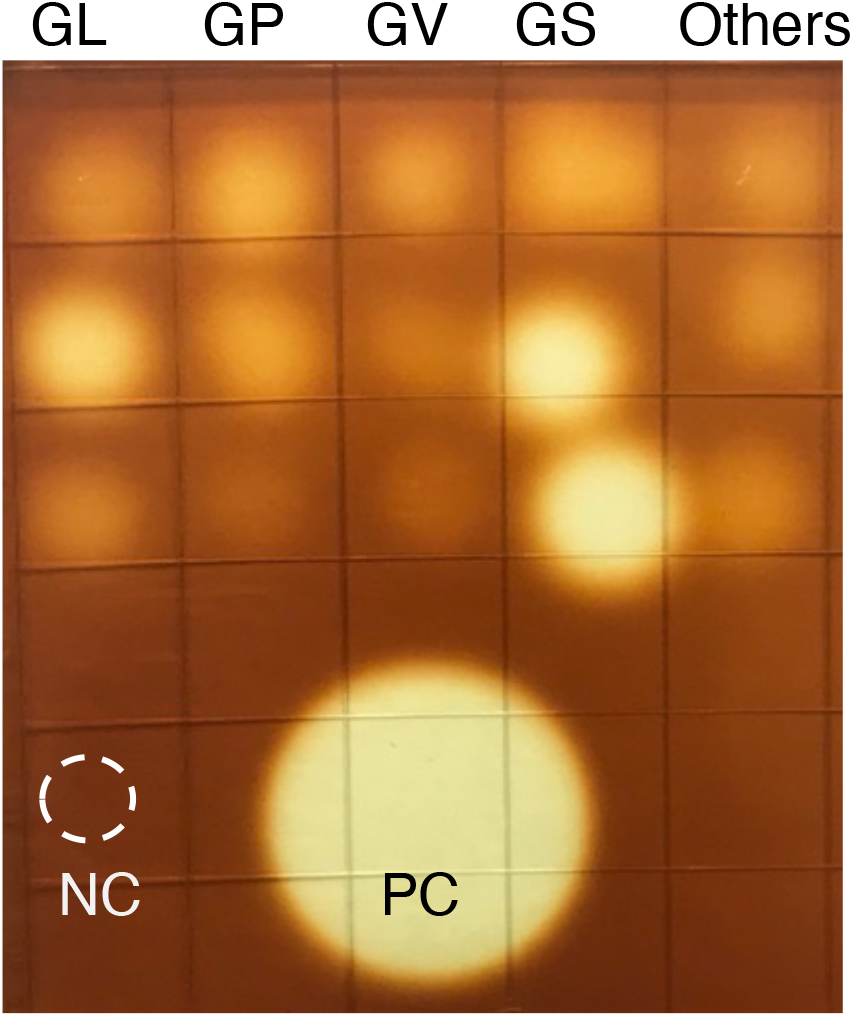
Amylase activity of cell free culture supernatants on glycogen agar. Each spot represents one isolate and a total of 15 different isolates, three each from GL: *G. leopoldii*, GP: *G. piotii*, GV: *G. vaginalis*, GS: *G. swidsinskii* and others [two isolates from *Gardnerella* genome sp. 3 (top & middle) and one isolate from unknown genome species (bottom) (corresponds to the subgroup D based on cpn60 classification system)] are shown. NC: mNYC, PC: *B. licheniformis* amylase (0.05 mg/ml).

**Figure 2:**
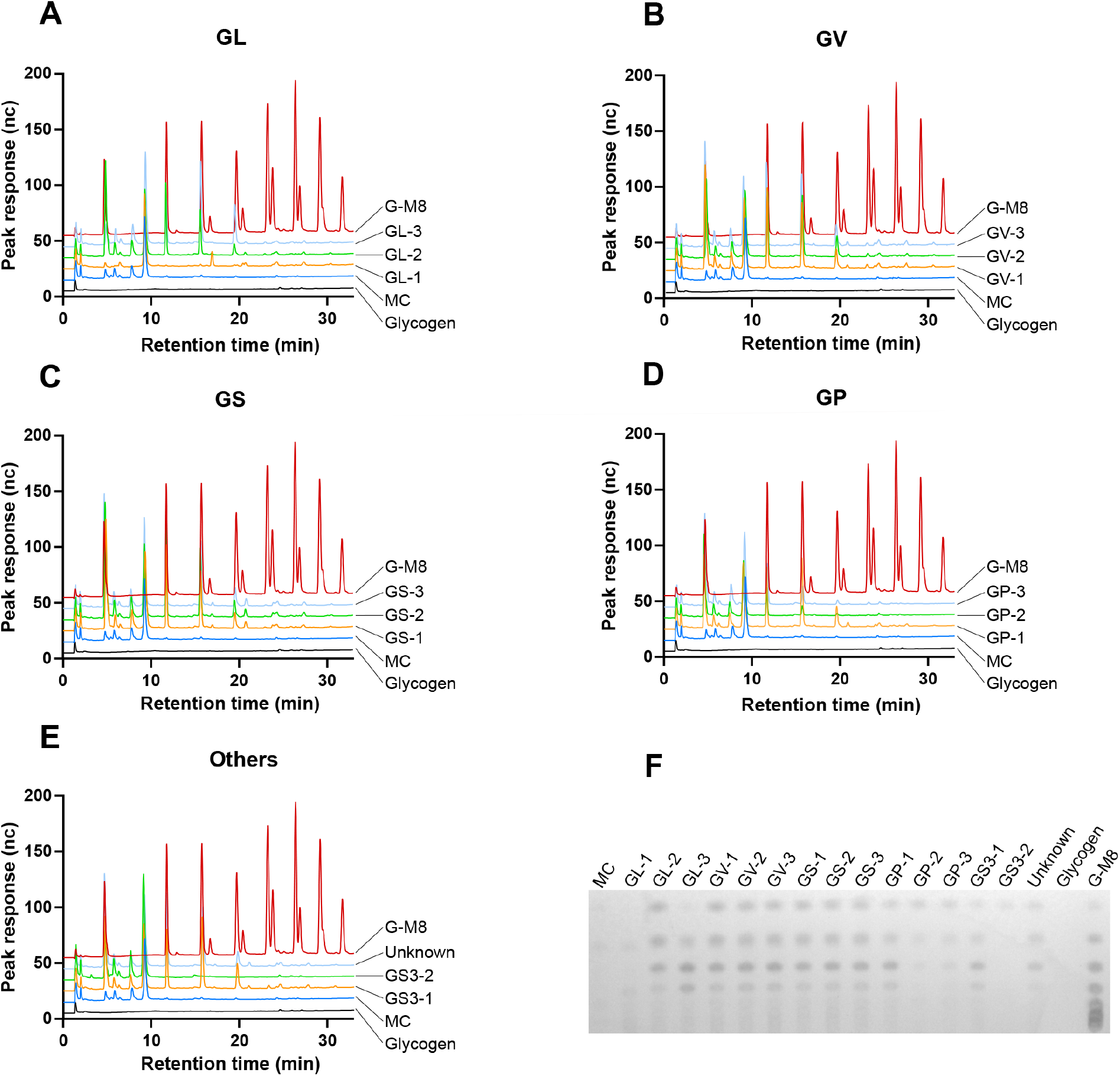
Identification of glycogen breakdown products produced after 24 h by culture supernatants of 15 *Gardnerella* isolates by HPAEC-PAD (A-E) and TLC (F). GL: *G. leopoldii*, GV: *G. vaginalis*, GS: *G. swidsinskii*, GP, *G. piotii*, and others [two isolates from *Gardnerella* genome sp. 3 (top & middle) and one isolate from unknown genome species (bottom) (corresponds to the subgroup D based on cpn60 classification system)], MC: media control, G-M8: glucose to maltooctaose standards.

### Identification of genes encoding carbohydrate active enzymes in *Gardnerella* genomes

Out of 20,017 predicted protein sequences from 15 isolates uploaded into the dbCAN2 webserver, 886 sequences had at least one carbohydrate active domain. Results were filtered to include only glycosyl hydrolase (GH) domains associated with a signal peptide. Only 52/836 sequences identified as belonging to the GH class had an associated signal peptide. Results were further filtered to identify predicted extracellular glycogen degrading enzymes (protein containing at least one GH13 domain associated with signal peptide), resulting in identification of 29 sequences.

A multidomain enzyme containing an N-terminal α-amylase domain (GH13_32), followed by 2 - 4 carbohydrate binding modules (CBM, of types CBM25, CBM41 or CBM48) and a C-terminal pullulanase domain (GH13_14) was identified in all isolates except N170. This enzyme also included an N-terminal signal peptide domain, and a potential sortase sorting signal including an LPxTG motif (Figure 3A). This multidomain enzyme is traditionally referred to as Type II amylopullulanase, however, recently it has been suggested that these enzymes are more appropriately classified as Type II α-amylase-pullulanase (23) and so the term ‘a-amylase-pullulanase’ is used throughout this paper. The domain organization of *G. leopoldii* NR017 α-amylase-pullulanase (RFT33565.1) is shown in Figure 3A. Similarly, a single α-amylase (GH13_32) domain containing α-amylase enzyme with an N-terminal signal peptide and a C-terminal transmembrane domain was identified in all 15 isolates (NR017 accession number RFT33564.1) (Figure 3B).

**Figure 3:**
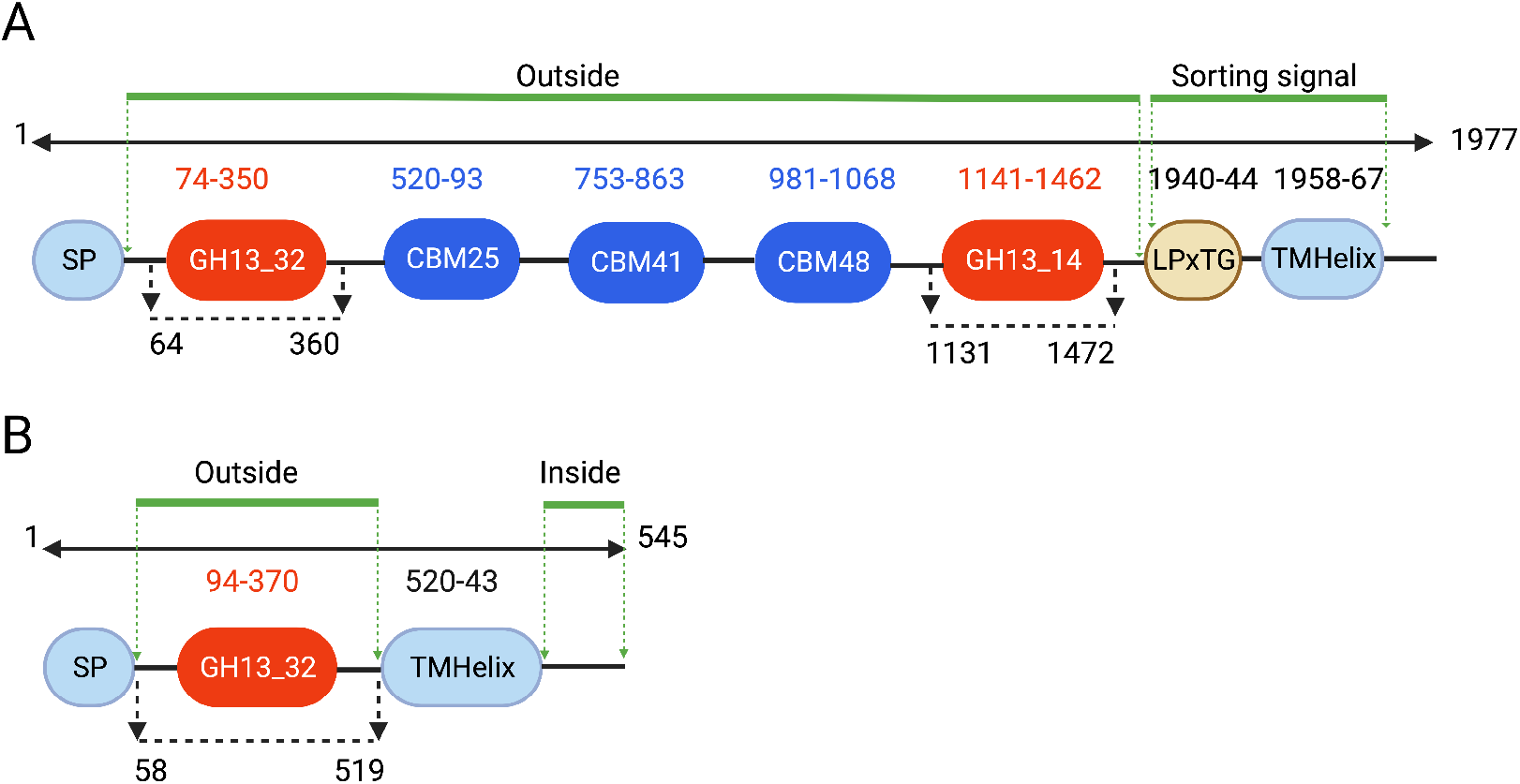
Predicted domains of *G. leopoldii* NR017 α-amylase-pullulanase (1977 amino acids) (A) and α-amylase (545 amino acids) (B). Amylase and pullulanase domains belonging to GH13 subfamilies 32 and 14 are indicated in red while three carbohydrate binding modules (CBM) are in blue. SP, Signal peptide; Sorting signal: LPxTG motif and a hydrophobic transmembrane helix domain (TMHelix) domain. Numbering indicates amino acid positions. The regions used for recombinant protein expression are indicated with black broken line with arrows. Predicted extracellular and intracellular regions are indicated with green lines.

### Phylogenetic analysis of GH13_32 amylase domains and relationship with other amylases

GH13_32 amylase domain sequences from the single domain α-amylase and α-amylase-pullulanase from all study isolates formed two distinct clusters (Figure 4A). Intra-cluster amino acid sequence identity of amylase domains within single domain α-amylase and α-amylase-pullulanase enzymes was 75-100% and 66-100%, respectively, whereas inter-cluster identity was 64-74%.

GH13 enzymes have a conserved catalytic triad of two aspartate residues (acts as a nucleophile and to stabilize the transition state) and a glutamate residue (general acid base) (24). *In silico* analysis of GH13_32 amylase domains of *G. leopoldii* NR017 with GH13_32 amylase domains of functionally characterized α-amylase and α-amylase-pullulanase revealed conserved catalytic residues (Figure 4B). The catalytic triad was conserved across all GH13_32 amylase domains from 15 *Gardnerella* isolates. (Supplementary Figure 1).

**Figure 4:**
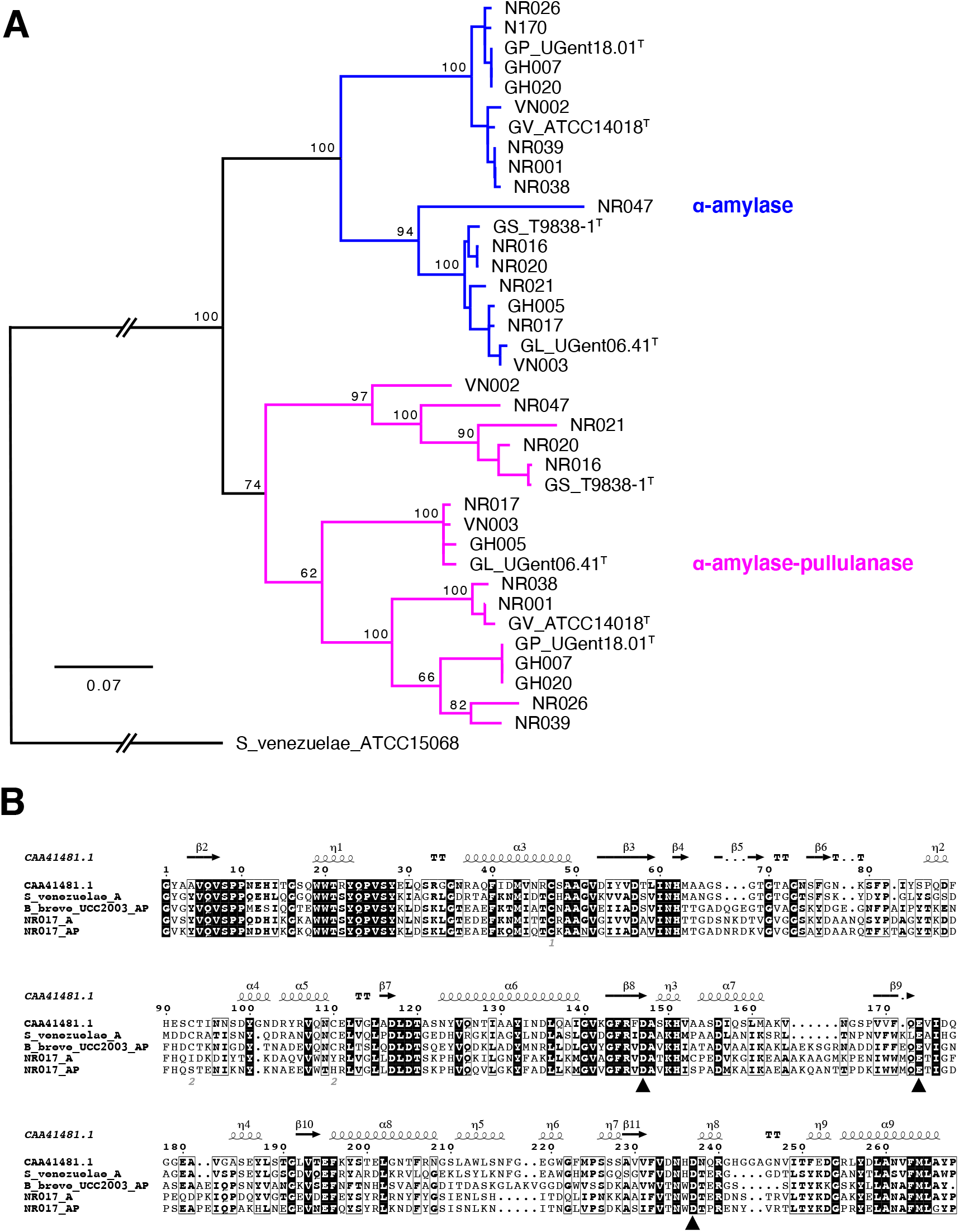
(A) Phylogenetic tree of all α-amylase domains from 15 *Gardnerella* isolates and 4 reference strains (indicated with T after the strain name). The tree is rooted with *Streptomyces venezuelae* ATCC15068. Trees are consensus trees of 100 bootstrap iterations and constructed using Neighbour joining method using Tamura-Nei distance model. The number at major branch points represents the percentage bootstrap support. (B) Sequence alignment of GH13_32 catalytic domains of *G leopoldii* NR017 α-amylase (NR017_A) and α-amylase-pullulanase (NR017_AP) with other functionally characterized GH13_32 members ((a-amylases from *Psuedoalternomonas haloplanktis* A23 (CAA41481.4) and *S. venezuelae* ATCC15068) and ‘amylopullulanase’ from *Bifidobacterium breve* UCC2003). Black triangles indicate conserved catalytic residues.

### Protein expression and purification

The entire open reading frame of *G. leopoldii* NR017 α-amylase-pullulanase (GH13_32 and GH13_14) comprises 5,933 bp encoding a 1,977 amino acid protein with a predicted mass of 215.3 kDa (Figure 3A). The gene segments encoding the α-amylase (amino acids 64-360) and pullulanase domains (amino acids 1131-1472) were each PCR amplified and ligated into pQE-80L vector to express as N-terminal 6His-tagged proteins in *E. coli* (Figure 5A, left panel). Expressed proteins were solubilized using 1% SDS and purified using nickel-nitriloacetic acid (Ni-NTA) columns (Supplementary Figure 2). Apparent molecular weights of the 6His-tagged amylase and pullulanase domains were 35.7 and 40.5 kDa, respectively (Figure 5B, left and middle).

**Figure 5:**
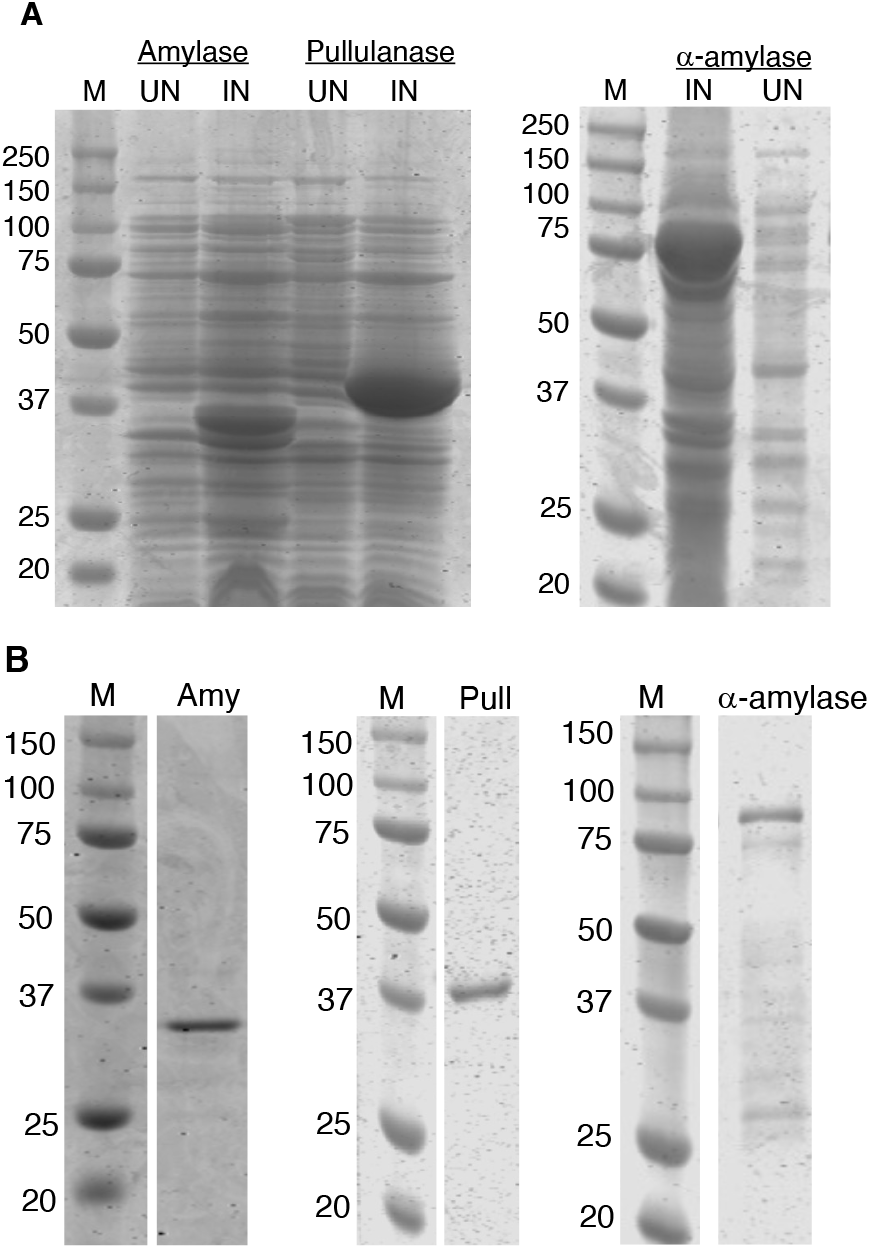
(A) Protein profiles of uninduced (UN) and induced (IN) *E. coli* cells containing pQE80L-AMY & pQE80L-PULL (top left) and pET41a+CO-AMY (top right) recombinant plasmids. (B) Purified 6His-tagged amylase (Amy) (bottom left) and pullulanase (Pull) (bottom middle) and GST-6His-tag α-amylase (bottom right). M molecular weight markers.

The *G. leopoldii* NR017 α-amylase enzyme (1635 bp encoding a 545 amino acid protein with a predicted mass of 60.5 kDa) has a predicted N-terminal signal peptide (amino acids 1-57) and C-terminal transmembrane domain (amino acids 520-545) (Figure 3B). A codon-optimized version of the truncated α-amylase gene sequence encoding amino acids 58-519 was expressed as a GST fusion protein with an apparent molecular weight of 79.9 kDa in *E. coli* BL21 DE3 cells (Figure 5A, right). The recombinant protein was solubilized using 1% SDS. Attempts to purify this GST fusion protein using glutathione beads resulted in a very low recovery of target protein due to poor binding to the beads (Supplementary Figure 3). The pET41a+amylase construct includes an N-terminal hexa-histidine tag between the GST tag and the amylase sequence. Purification of the fusion protein using a Ni-NTA column resulted in partial protein purification (Figure 5B, right). Additional lower molecular weight peptides observed in the column eluate were identified as degraded peptide fragments of the fusion protein by western blot with anti-His antibodies (Supplementary Figure 4).

### Activity of the purified proteins

The α-amylase domain from the α-amylase-pullulanase released maltose and maltotriose from maltopentose while maltose, maltotriose and maltotetraose were identified as the major products from maltodextrin (dextrose equivalent 13-17) and glycogen (Figure 6A). A small peak corresponding to isomaltose was observed in the HPAEC-PAD results from 24 and 48 h digests of glycogen with α-amylase domain (Figure 6A-middle right). The pullulanase domain produced maltotriose as the only product from pullulan (Figure 6B) while maltose was produced from glycogen (Figure 6C). Combined activities of the α-amylase and pullulanase domains on glycogen released maltose and maltotriose (Figure 6C) with no detectable maltotetraose. This suggests that the pullulanase domain can hydrolyze maltotetraose to release maltose but is less active (or inactive) on the smaller maltotriose substrate, perhaps because it is a poorer fit for the active site. To test this, the pullulanase domain was incubated with maltotriose, maltotetraose and maltopentose. The results showed that pullulanase hydrolyzed maltopentose (into maltose and maltotriose), and to a lesser extent, maltotetraose (into maltose) and maltotriose (into glucose and maltose) (Supplementary Figure 6). Overall, our results show that α-amylase and pullulanase domains from α-amylase-pullulanase can hydrolyse a-1,4, and a-1,4 and a-1,6 glycosidic bonds in glycogen, respectively.

**Figure 6.**
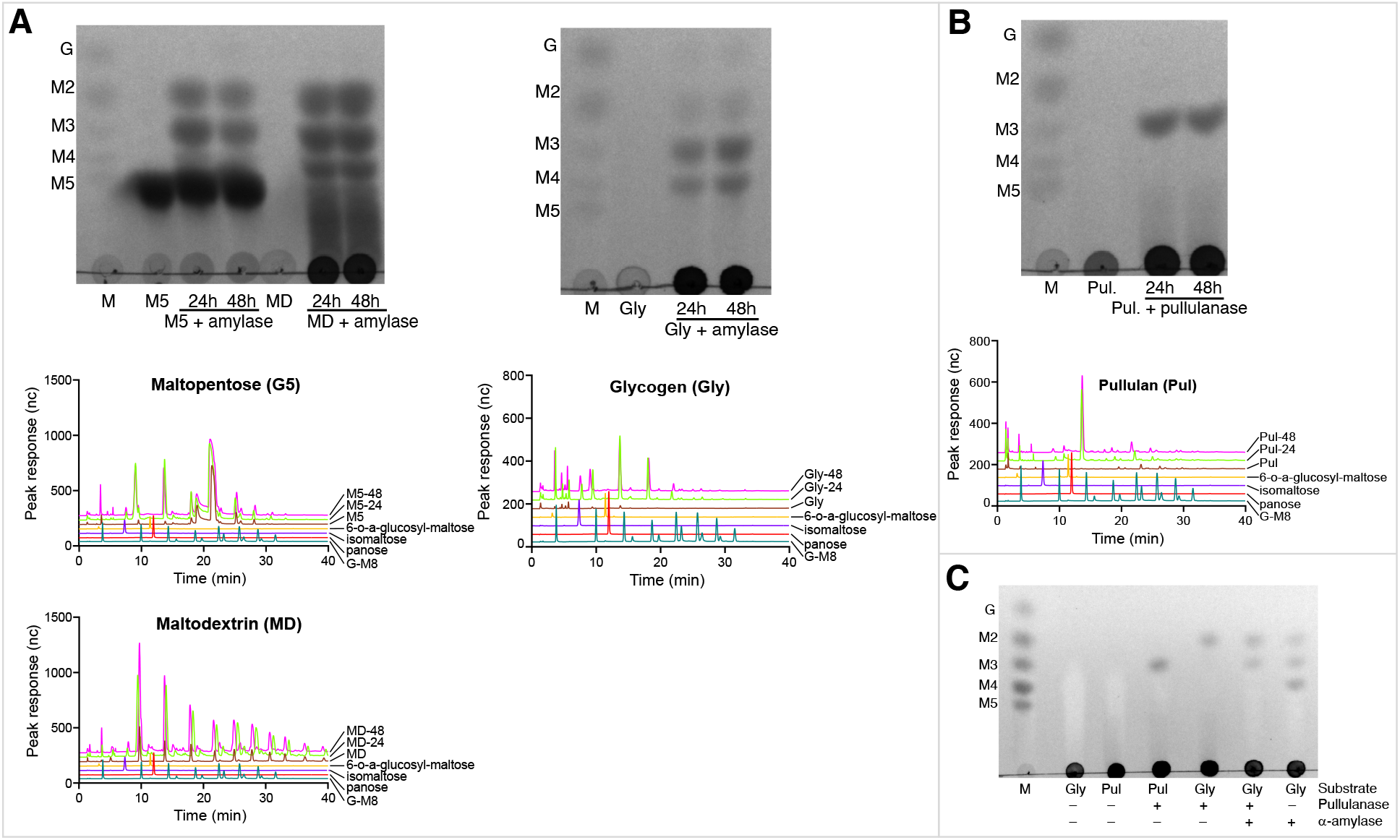
Identification of products released by α-amylase domain (GH13_32) of α-amylase-pullulanase from maltopentose (M5) and maltodextrin (MD) and glycogen (Gly) after 24 and 48h of incubation using TLC (top) and HPAEC-PAD (middle and bottom). (B) Products released from pullulan by the pullulanase domain (GH13_14) after 24 and 48 h incubation were identified using TLC (top) and HPAEC-PAD (bottom). (C) Identification of products released after 24h from pullulan or glycogen by pullulanase, α-amylase or a mixture of α-amylase and pullulanase domains. Standards in Lane M of each panel: G: glucose, M2: maltose, M3: maltotriose, M4: maltotetraose and M5: maltopentose.

Since the α-amylase domain from the single-domain α-amylase enzyme was expressed as a GST-6xHis fusion, we initially confirmed that no amylase activity was detected from the GST-6xHis tag (Supplementary Figure 5). Maltose and maltotriose were produced when maltopentose was incubated with single domain α-amylase (Figure 7A, top and middle). Similarly, maltose, maltotriose and maltotetraose were identified as the major products in the reaction mixture of glycogen with single domain α-amylase (Figure 7B). As expected, the maltodextrin substrate included maltose, maltotriose and maltotetraose, however, an increase in intensity of maltose, maltotriose and maltotetraose spots was observed in TLC results in the reaction mixture of single domain α-amylase with maltodextrin when compared to a substrate only control (Figure 7A, top). HPAEC-PAD analysis of products from maltodextrin also showed larger peaks of maltose, maltotriose and maltotetraose relative to the control (Figure 7A, bottom).

**Figure 7:**
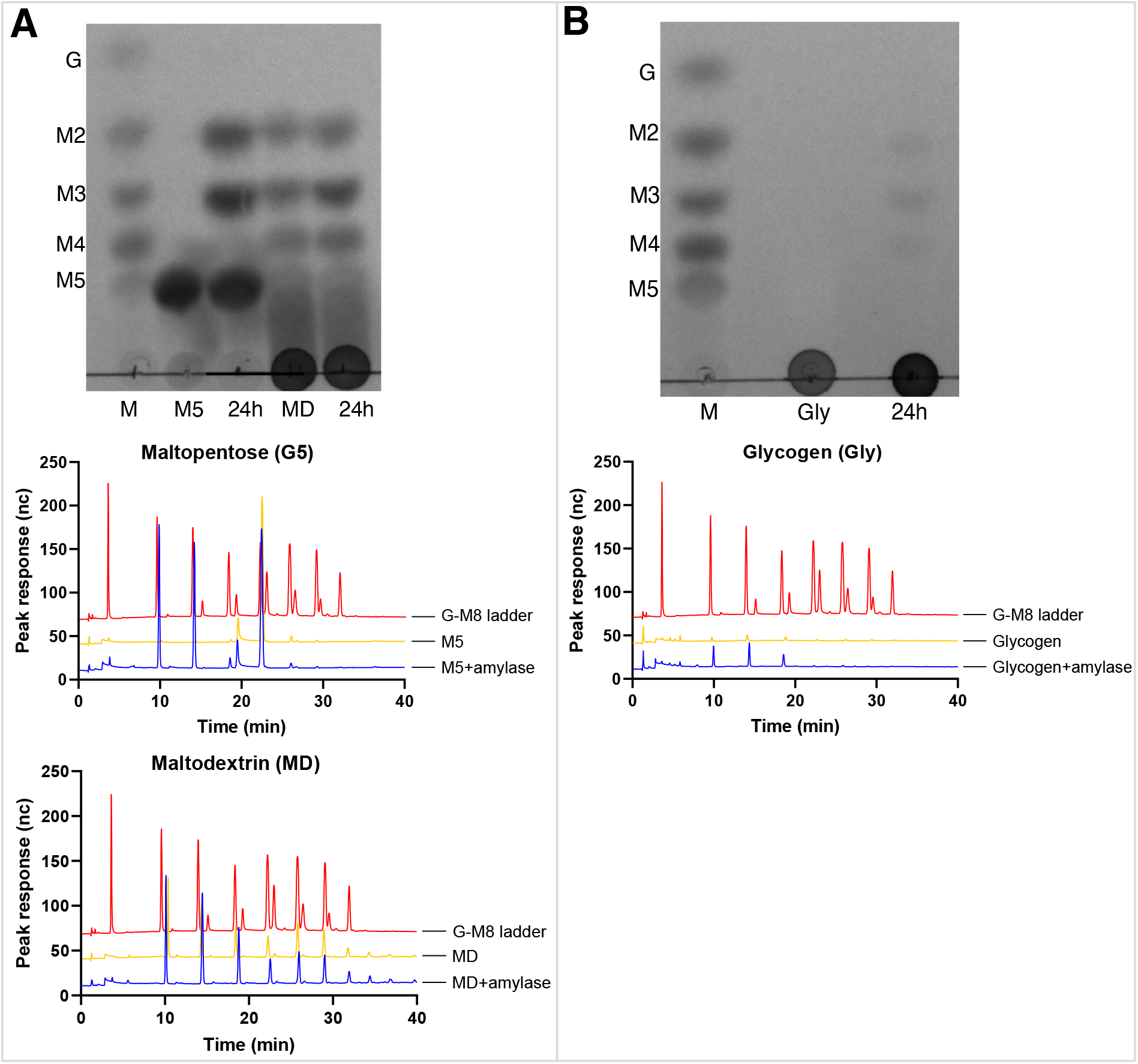
(A) Identification of products released by the purified single domain α-amylase enzyme from maltopentose (M5) and maltodextrin (MD) using TLC (top) and HPAEC-PAD (bottom) after 24h incubation. (B) Products released from glycogen by the purified single domain α-amylase enzyme after 24 h incubation were identified using TLC (top) and HPAEC-PAD (bottom). Standards in Lane M of each panel: G: glucose, M2: maltose, M3: maltotriose, M4: maltotetraose and M5: maltopentose.

## Discussion

Dysbiosis in the vaginal microbiome has significant consequences that can affect the quality of life and reproductive health of women. There are several hypotheses about how dysbiosis is initiated and maintained (25, 26) but the ecological factors determining microbial population dynamics in the vaginal microbiome are not well understood. In addition to direct antagonistic (27) or synergistic (28) interactions among microbiota, changes in environmental conditions due to pregnancy status, hormone levels, infection, sexual behaviour and hygiene practices certainly play a role in determining the fitness of vaginal microbiota and thus microbial community composition. *Gardnerella* spp. have been implicated in the initial stages of transition to bacterial vaginosis (26) and so factors affecting their fitness, including their interaction with vaginal glycogen, are of particular interest.

Glycogen breakdown is a complex and multistep process that requires the collective action of several glycogen degrading enzymes. Vaginal secretions contain host derived pancreatic α-amylase and α-glucosidases that can digest glycogen into smaller sugars (13). Until recently, the contribution of bacterial enzymes in this process was unclear. Recently, van der Veer *et al*. showed that vaginal isolates of *Lactobacillus crispatus* are capable of growing in glycogen supplemented medium and encode type I pullulanase in their genomes (29). This study further reported that strains showing less efficient or no growth on glycogen have an N-terminal deletion mutation in their putative type I pullulanase gene suggesting its role in extracellular glycogen digestion. Forney *et al*. identified eight putative amylases from vaginal metagenomes of *Bifidobacterium vaginale, B. lacrimalis, L. gasseri, L. crispatus, L. iners* and *L. jensenni* and further reported that multiple microbial amylases are present in vaginal fluid (14). Similarly, Woolston *et al*. identified putative glycogen degrading enzymes in genomes of vaginal bacteria including *Gardnerella* spp. and biochemically characterized the glycogen degradation ability of six GH13 hydrolase enzymes (30) identified in those species. Together these findings suggest that bacterial enzymes contribute to the breakdown of vaginal glycogen.

The genus *Gardnerella* was a mono-specific genus until recently when it was classified into four species and additional “genome species” that have not yet been named due to insufficient evidence (4). The role of *Gardnerella* spp. in extracellular glycogen digestion and whether the species in this genus differ in digesting glycogen is not clear. Although earlier studies reported that *Gardnerella* isolates can utilize glycogen (31–33), it is not clear if all newly defined species are capable of digesting extracellular glycogen. In this study, we used cell free culture supernatants from genetically characterized isolates to test extracellular glycogen breakdown ability of different *Gardnerella* spp.

Selection of an appropriate culture medium to make culture supernatants is a challenge. As our main interest was to identify the sugars released from glycogen, differentiating the background sugars that are already present in medium from those obtained from glycogen hydrolysis is crucial. *Gardnerella* is typically cultured in BHI broth or NYCIII and both media contain glucose. In addition, the complement inactivated horse serum (obtained by heat treatment at 56°C for 20 minutes) added to NYCIII can itself digest glycogen. It is reported that heat treatment at 70°C for 20 minutes inhibits the maltase and amylase activity of fetal bovine serum but not horse serum (34). For this reason, in our experiments a modified NYCIII media supplemented with heat inactivated (70°C for 20 minutes) fetal bovine serum instead of horse serum and no added glucose was used to prepare culture supernatants. Cell free culture supernatants of all isolates tested showed amylase activity in a glycogen agar plate assay, suggesting that glycogen degradation is a conserved property in the genus *Gardnerella*. Glucose, maltose, maltotriose and maltotetraose were identified as the major products released from glycogen by culture supernatants of all *Gardnerella* isolates except N170 isolate which produced only glucose. Consistent with this observation α-amylase-pullulanase genes were detected in all isolates except N170. Overall, no species-specific patterns of glycogen breakdown products were found in *Gardnerella* spp.

Of the two extracellular glycosyl hydrolases identified by dbCAN2, the α-amylase-pullulanase was predicted to contain separate amylase and pullulanase catalytic domains; a conformation historically referred to as Type II amylopullulanase (35). With the availability of more sequence data, these enzymes are now classified as Type II α-amylase-pullulanase and they differ from amylopullulanases which have only one catalytic domain capable of hydrolyzing both a-1,4 and a-1,6 glycosidic bonds (23). The amylase and pullulanase domains of the identified *Gardnerella* α-amylase-pullulanase flanked 2 - 4 CBMs. CBMs appended to GH13 enzymes are important for binding to different a-glucans, such as amylose, amylopectin, glycogen, pullulan and oligosaccharides derived from these polysaccharides (36, 37). CBMs increase the concentration of enzyme on the substrate surface and their removal can significantly reduce enzyme activity. It is reported that removal of CBMs reduced the activity of *Eubacterium rectale* α-amylase on starch by ~40 fold compared to the wildtype protein (38). Similarly, compared to a full-length enzyme, removal of CBM domains reduced the activity of *B. adolescentis* P2P3 Type II pullulanase on glycogen (39). We did not investigate the roles of the CBMs in the α-amylase-pullulanase characterized in this study since our attempts to express full-length α-amylase-pullulanase were unsuccessful.

Recently, Woolston *et al*. (30) characterized a full length ‘amylopullulanase’ (2013 amino acids) from a vaginal isolate of *G. vaginalis* and reported maltose and maltotriose as the major products from glycogen. In this report, the authors described the enzyme as containing ‘a-amylase domains’ rather than amylase and pullulanase catalytic domains as we describe here. By expressing the two catalytic domains separately we have demonstrated that they have distinct activities, hydrolysing a-1,4, and a-1,4 and a-1,6 bonds, respectively.

Pullulanase degrading enzymes are classified into two major types, pullulanases (type I and type II) and pullulanase hydrolases (type I, II and III) based on substrate specificities and reaction products. While type I pullulanases hydrolyze only a-1,6 glycosidic bonds, type II pullulanases are capable of hydrolyzing a-1,4 and a-1,6 glycosidic bonds. Amylopullulanase, a subtype of type II pullulanase, are mainly reported in thermophilic bacteria and have a single catalytic domain, whereas α-amylase-pullulanase has dual catalytic domains (a-amylase: for a-1,4 glycosidic bonds and pullulanase: for a-1,6 glycosidic bonds) and are reported in mesophilic bacteria (23). α-amylase-pullulanases have been characterized in *Bifidobacterium breve* UCC2003, *B. adolescentis* P2P3 (39, 41) and a few other species (23). Our results showed that the pullulanase domain of the *G. leopoldii* α-amylase-pullulanase hydrolyzes a-1,4 as well as a-1,6 glycosidic bonds, which to our knowledge has not been reported before. Kim *et al*. (2021) reported that the pullulanase domain of an α-amylase-pullulanase from *B. adolescentis* P2P3 is capable of hydrolyzing a-1,6 glycosidic bonds and releasing maltotriose from pullulan, however no products were detected with amylose indicating a lack of activity on a-1,4 linkages (39). The demonstrated activity of the *G. leopoldii* pullulanase domain on a-1,4, and a-1,6 glycosidic bonds would explain why maltose was the only detectable product released from glycogen (Figure 6C) and suggests that this domain might better be described as an amylopullulanase. It is important to note, however, that the lack of CBMs associated with the domains we investigated in the current study could affect their activity and the range of products released from different substrates and so further study of the influence of CBMs is warranted.

Both α-amylase and α-amylase-pullulanase enzymes are active against oligosaccharides and glycogen. Although the overall sequence identity between the two amylase domains characterized in this study was only 64-74%, predicted catalytic residues were perfectly conserved (Figure 4B). Either domain would thus be capable of acting on the malto-oligosaccharides produced by the debranching activity of the pullulanase domain to generate glucose, maltose, and other small oligosaccharides. This type of functional redundancy where two enzymes have similar biochemical activities is common in bacteria (40).

The *Gardnerella* α-amylase and α-amylase-pullulanase are predicted to be extracellular based on the presence of N-terminal signal peptides. The α-amylase contains a C-terminal transmembrane helix, suggesting that the protein is inserted in the cell membrane. In the α-amylase-pullulanase sequence an LPxTG motif was identified immediately N-terminal to the predicted transmembrane helix (Figure 3A), suggesting this protein is anchored to the cell wall through the sortase mechanism. The presence of these features suggests that both enzymes are tethered to the cell envelope. Cell envelope anchored proteins can be released into the extracellular environment during bacterial growth as a by-product of cell wall turnover or cell death or proteolytic cleavage (42), which is consistent with our detection of amylase activity in sterile culture supernatants. Some *Gardnerella* species also produce cell wall anchored sialidases, NanH2 and NanH3, that are also released into the supernatant after cell death or by proteolytic cleavage (43).

Although glucose was identified as one of the major products in glycogen digests of *Gardnerella* culture supernatants, the purified α-amylase domain and single domain α-amylase proteins did not produce glucose from glycogen. TLC and HPAEC-PAD analysis showed that glucose was neither produced by the activity of culture media itself nor present in glycogen substrate (Figure 2F & 2G). Another possibility is that the activity of a full-length α-amylase-pullulanase protein on glycogen might generate different product profiles than the isolated domains. A full-length type II pullulanase enzyme (containing two separate α-amylase and pullulanase domains) from *Bifidobacterium adolescentis* P2P3 was reported to produce maltose only from glycogen, while the purified amylase domain protein from the same enzyme produced a mixture of glucose, maltose and maltotriose (39). Glucose may also be produced by the activity of intracellular a-glucosidase on malto-oligosaccharides generated from glycogen hydrolysis that are released by dead/lysed bacteria in the culture. We have previously shown that *Gardnerella* spp. have conserved intracellular a-glucosidase that can release glucose from smaller oligosaccharides (44).

Our findings show that all *Gardnerella* species produce extracellular α-amylase and α-amylase-pullulanase enzymes that are ‘public goods’ in the vaginal microbiome, breaking down glycogen to produce similar fingerprints of glycogen derived products and contributing to the nutrient pool available to the vaginal microbiota (Figure 8). Determining if these *Gardnerella* spp. differentially utilize and compete for these breakdown products will provide insights into whether competition for malto-oligosaccharides plays a role in determining *Gardnerella* community structure in the vaginal microbiome.

**Figure 8:**
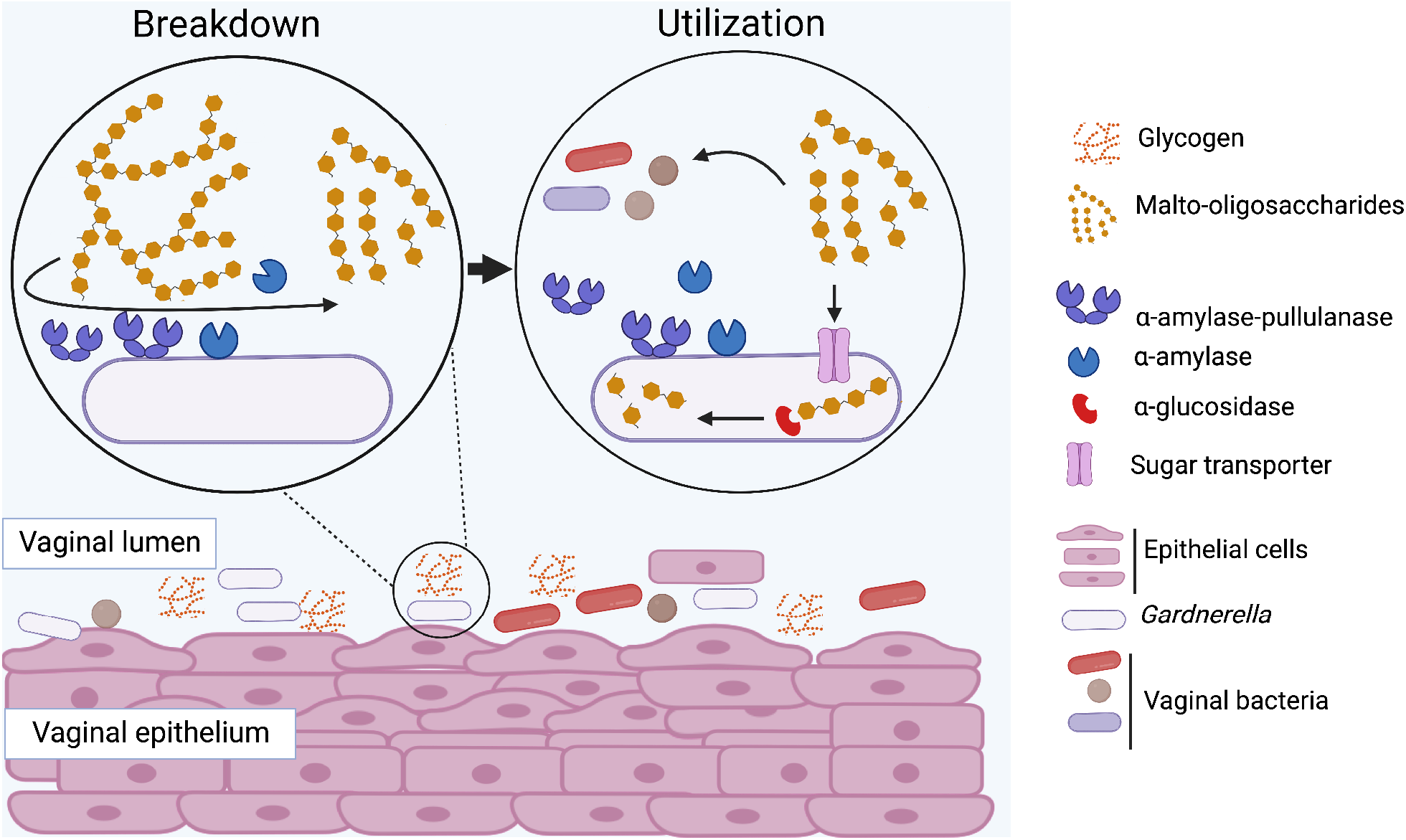
Proposed mechanisms of extracellular glycogen utilization in *Gardnerella* spp. Glycogen is digested into malto-oligosaccharides by α-amylase-pullulanase and α-amylase and these breakdown products are transported inside and can be further digested to glucose by other glycosyl hydrolases including a previously characterized a-glucosidase enzyme (44). Extracellular glycogen breakdown products can also be utilized by other resident microbiota.

## Acknowledgements

This research was supported by a Natural Sciences and Engineering Research Council of Canada Discovery Grant (J.E.H.) and Agriculture and Agri-Food Canada (D.W.A. project no. J-002262). P.B. is supported by a Devolved Scholarship from the University of Saskatchewan. The authors would like to thank Champika Fernando for technical support, Shubham Dutta for guidance with Western blot, and Noreen Rapin and Dr. Tony Ruzzini for providing reagents.

