## Supplementary Figures for "Glycogen degrading activities of catalytic domains of α-amylase and α-amylase-pullulanase enzymes conserved in *Gardnerella* spp. from the vaginal microbiome"

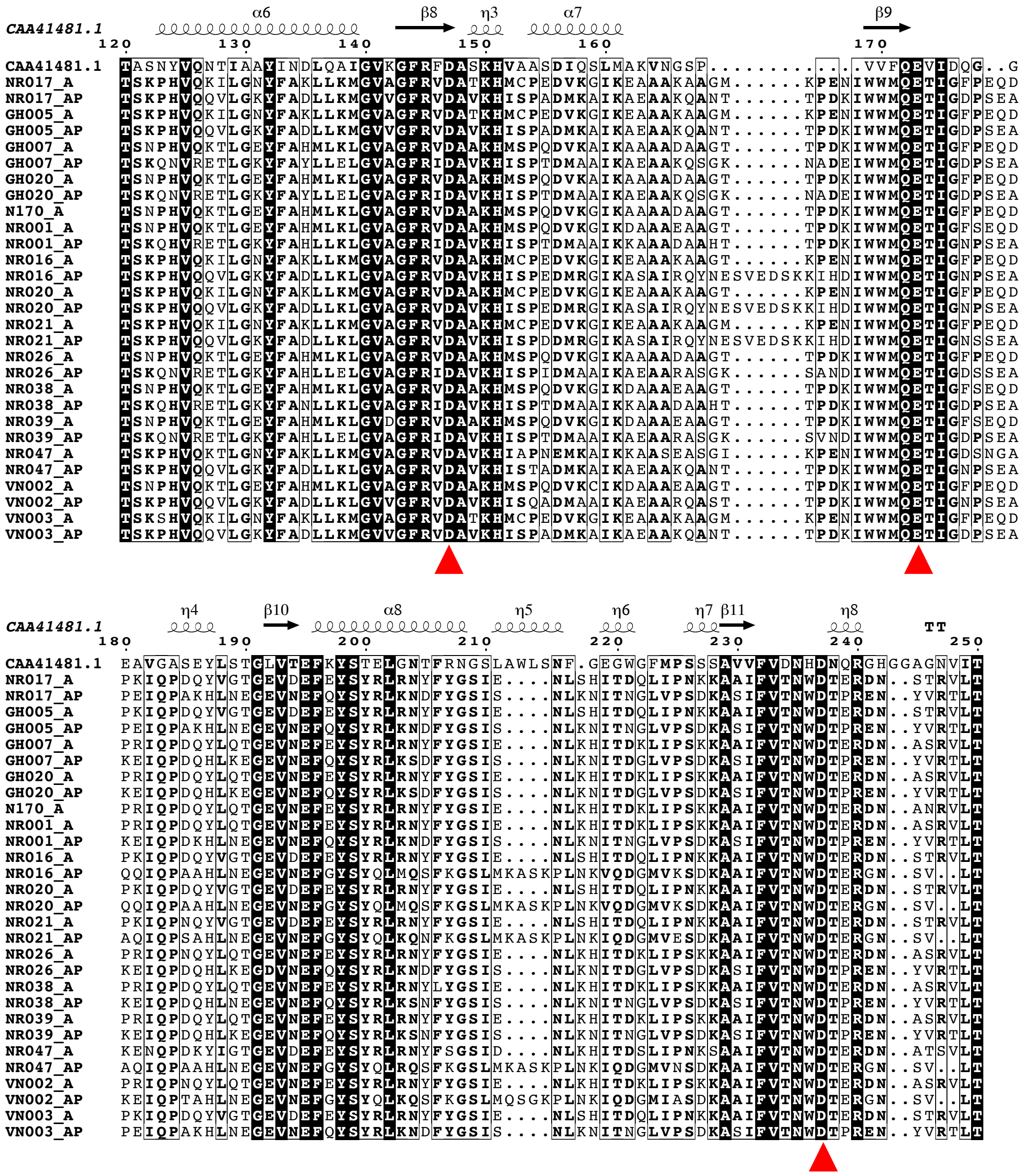

Supplementary Figure 1: Sequence alignment of all GH13_32 domains from 15 *Gardnerella* isolates with GH13_32 domain of functionally characterized α-amylase from *P. haloplanktis* A23 (CAA41481.1). An alignment including amino acids from 120 to 250 positions only is shown. Conserved catalytic residues are indicated with red triangles and invariant sequences are highlighted in dark background.

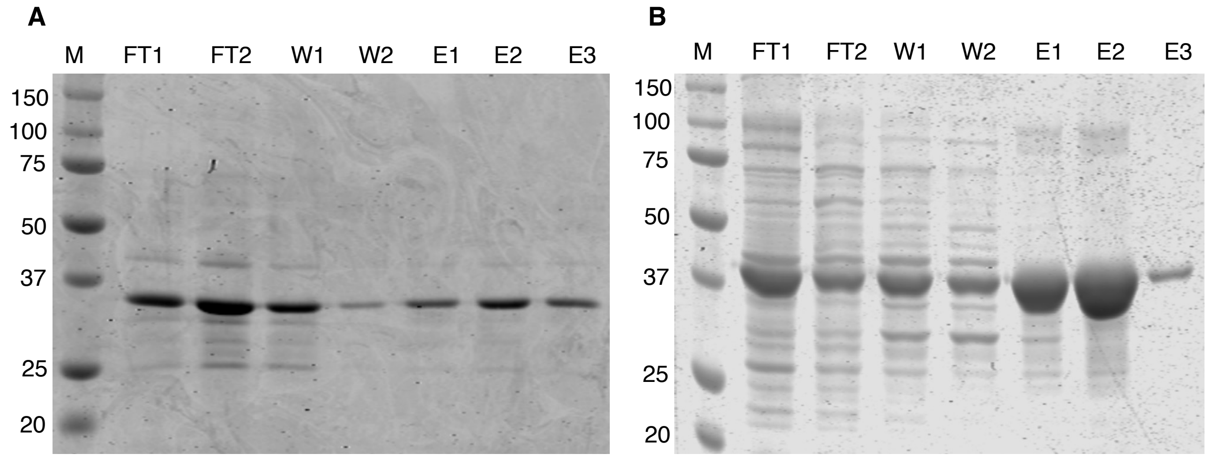


Supplementary Figure 2: Ni-NTA purification of 6His-tagged amylase (A) and pullulanase (B) domain proteins. M: molecular weight marker, FT1 &FT2: flow through fractions, W1&W2: wash fractions, E1-E3: elution fractions.

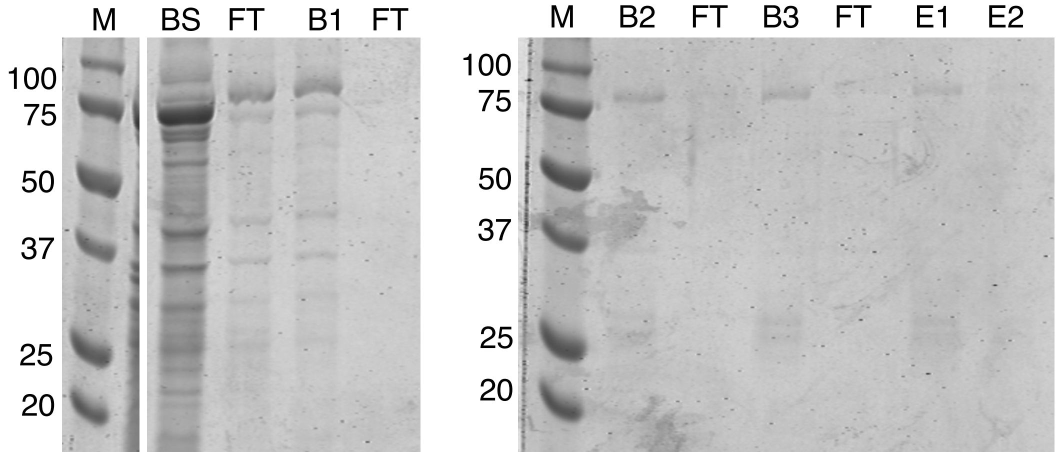


Supplementary Figure 3: Analysis of fractions obtained during protein purification using glutathione beads. M: molecular weight markers, BS: mixture of glutathione beads and solubilized protein, FT: flowthroughs, B1-B3: Protein bound glutathione beads obtained after first, second and third wash, E1-E2: elution fractions

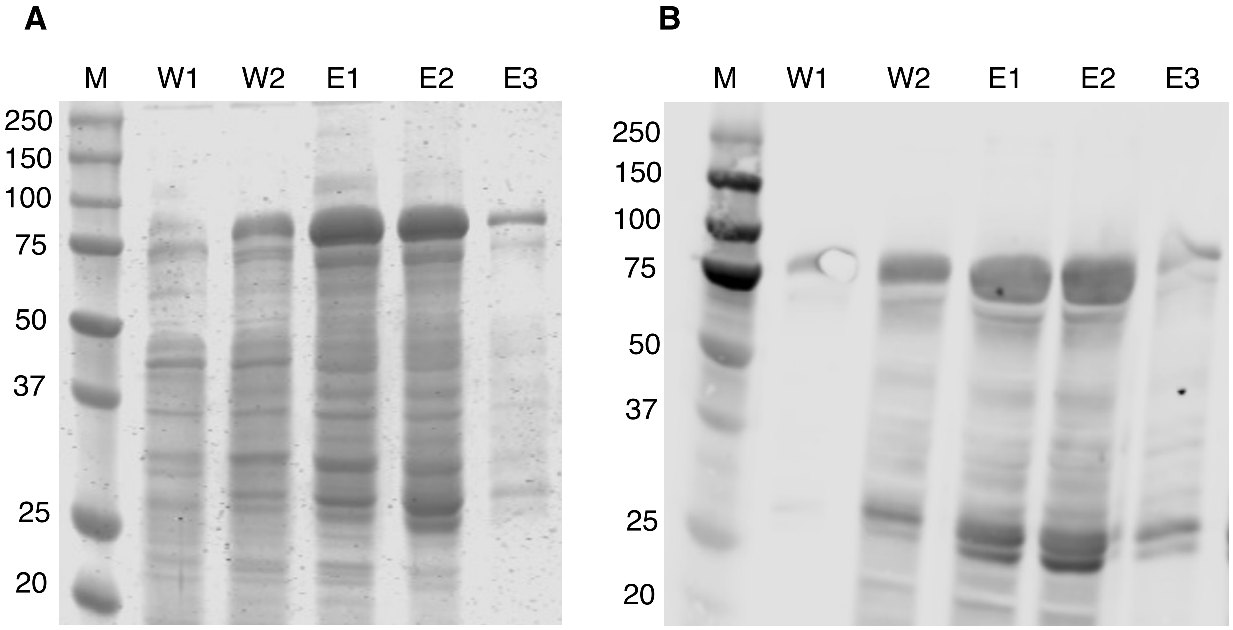


Supplementary Figure 4: SDS-PAGE (A) and Western blot (B) analysis of fractions obtained during recombinant α-amylase protein purification using Nickel affinity chromatography (A). Immunoblot membrane was first incubated with primary antibody (Anti-6x His tag monoclonal antibody, 1:2,000 dilutions) followed by washing and further incubation with secondary antibody conjugates (Alexa fluor 680 goat anti-mouse IgG, 1: 20,000 dilutions). Finally, membrane was scanned at 700 nm using Li-COR scanner after secondary antibody wash. M- molecular weight markers, W1 & W2: wash fractions, E1-E3: elution fractions.

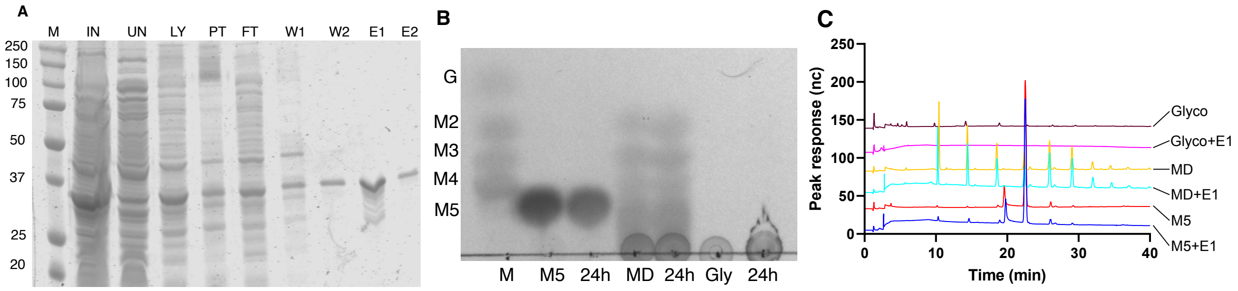


Supplementary Figure 5: (A) Expression and Ni-NTA purification of GST-His fusion tag. M: molecular weight markers, IN: IPTG induced culture, UN: uninduced culture, LY: lysate, PT: pellet, FT: flowthrough, W1 and W2: wash fractions, E1 and E2: elution fractions. Identification of products obtained from purified GST-His tag with maltopentose (M5), maltodextrin (MD) and Glycogen (Gly) using TLC (B) and HPAEC-PAD (C). M: sugar standards: G: glucose, M2: maltose, M3: maltotriose, M4: maltotetraose.

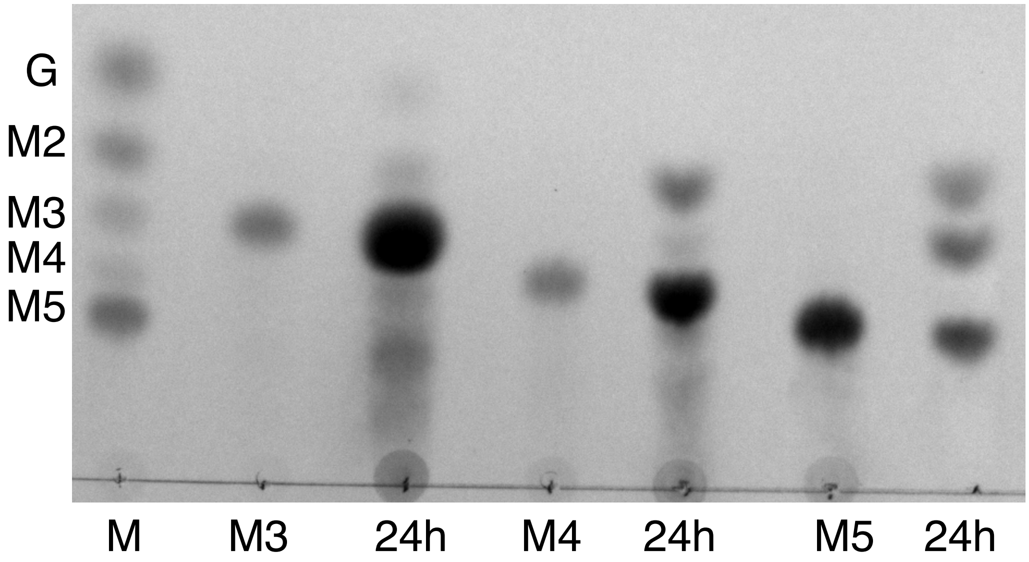


Supplementary Figure 6: Identification of products obtained from purified pullulanase domain protein with maltotriose (M3), maltotetraose (M4) and maltopentose (M5) after 24h of incubation. M: sugar standards: G: glucose, M2: maltose, M3: maltotriose, M4: maltotetraose and M5: maltopentose.
